# Robustness of the autophagy pathway to somatic copy number losses

**DOI:** 10.1101/2022.04.29.489531

**Authors:** Pierfrancesco Polo, Niklas Gremke, Thorsten Stiewe, Michael Wanzel

**Affiliations:** Institute of Molecular Oncology, Philipps-University, Marburg, Germany; Department of Gynecology and Obstetrics, Philipps-University Marburg, Germany; Institute of Lung Health (ILH), Giessen, Germany; Universities of Giessen and Marburg Lung Center, German Center for Lung Research (DZL), Marburg, Germany

**Keywords:** autophagy, cancer metabolism, aerobic glycolysis, somatic copy number alterations, SCNA, copy number loss, haploinsufficiency, metabolic cancer therapy

## Abstract

Autophagy allows cells to temporarily tolerate energy stress by replenishing critical metabolites through self-digestion, thereby attenuating the cytotoxic effects of anticancer drugs that target tumor metabolism. Autophagy defects could therefore mark a metabolically vulnerable cancer state and open a therapeutic window. While mutations of autophagy genes (ATGs) are notably rare in cancer, haploinsufficiency network analyses across many cancers have shown that the autophagy pathway is frequently hit by somatic copy number losses of ATGs like *MAP1LC3B/ATG8F* (*LC3*), *BECN1/ATG6* (Beclin-1), and *ATG10*. Here, we used CRISPR/Cas9 technology to delete increasing numbers of copies of one or more of these ATGs in non-small cell lung cancer cells and examined the effects on sensitivity to compounds targeting aerobic glycolysis, a hallmark of cancer metabolism. Whereas complete knock-out of one ATG blocked autophagy and led to profound metabolic vulnerability, this was not the case for combinations of different non-homozygous deletions. In cancer patients, the effect of ATG copy number loss was blunted at the protein level and did not lead to accumulation of p62 as a sign of reduced autophagic flux. Thus, the autophagy pathway is shown to be markedly robust and resilient, even with concomitant copy number loss of key autophagy genes.

## 1. Introduction

Cancer cells reprogram their energy metabolism to produce the enormous amounts of biomass needed for sustained tumor growth [1]. Already Otto Warburg noted a high glycolytic activity of tumor cells even in the presence of oxygen (aerobic glycolysis). Today, this characteristic metabolic phenotype (called ‘Warburg effect’) is considered a hallmark of cancer cells and attributed to a special dependence of highly proliferating cells on biosynthetic pathways that use intermediates derived from glycolysis [2-4]. At the molecular level, this is achieved by increased uptake of glucose and downstream inhibition of glycolysis at the level of pyruvate kinase and pyruvate dehydrogenase. These processes can be pharmacologically targeted with 2-deoxy-D-glucose (2DG) and dichloroacetate (DCA), respectively [5]. 2DG is a glucose analog that is taken up by cells through glucose transporters and phosphorylated by hexokinase. It cannot be further metabolized in the glycolysis pathway and therefore considered a glycolysis inhibitor. DCA, a structural analog of pyruvate, functions as an inhibitor of pyruvate dehydrogenase kinase 1 (PDK1), thus increases pyruvate dehydrogenase activity and stimulates efflux from glycolysis into the tricarboxylic acid cycle [6]. Although the impact of 2DG and DCA on cancer cell metabolism is far from being understood, both compounds are widely considered as anti-Warburg drugs that interfere with the central role of glycolysis in cancer cells [5]. Studies using 2DG and DCA have provided deeper mechanistic insight into cancer cell metabolism and have led to the development of novel therapeutic strategies to exploit the altered metabolism [7]. In clinical trials 2DG and DCA showed dose-limiting toxicity preventing their clinical use [8]. Some newer drugs are currently being evaluated in preclinical models or clinical trials [1, 8], but it appears that many are still limited more by toxicities than by their ability to kill cancer cells [7]. A better understanding of the metabolic dependencies in specific tumor tissues holds the key for defining the aspects of metabolism most limiting for tumor growth and finding a therapeutic window to exploit those vulnerabilities for better cancer treatment [7].

Macro-autophagy (hereafter referred to as autophagy) is a physiological and evolutionary conserved self-digestion process, in which intracellular components are sequestered in double-membrane vesicles called autophagosomes that fuse with lysosomes for recycling of nutrients [9-11]. Its progression involves the core autophagy machinery, encoded by autophagy-related genes (ATG), and can be subdivided into the sequential steps: induction, nucleation, elongation, docking and fusion, degradation and recycling [12]. Induction is inversely controlled by the mammalian target of rapamycin (mTOR) and the 5’ adenosine monophosphate-activated protein kinase (AMPK). Whereas mTOR negatively regulates autophagy [13], the cellular energy sensor AMPK activates autophagy in response to various cellular stresses, including glucose starvation and treatment with inhibitors of aerobic glycolysis [14-18]. Both protein kinases phosphorylate, with opposite effects, the ULK1/2 kinase, that forms the pre-initiation complex together with FIP200, ATG13, ATG17, and ATG101 [19]. This, in turn, activates the Beclin-1 (also known as BECN1 or ATG6) initiation complex, which includes ATG4L, VPS34, UVRAG, and Bif-1. The Beclin-1 complex is essential for the nucleation of the phagophore, the immature form of the autophagosome, while vesicle elongation involves the microtubule-associated proteins 1A/1B light chain 3B (MAP1LC3B, also known as ATG8F or LC3), a member of the ATG8 protein family [20]. Newly synthesized pro-LC3 is processed by the ATG4 protease to LC3-I, and subsequently conjugated to phosphatidylethanolamine (PE) in a multistep process driven by the E1-like enzyme ATG7 and the E2-like enzymes ATG3 and ATG10 [21]. PE-conjugated (lipidated) LC3, commonly referred to as LC3-II, is anchored to the phagophore membrane and functions in the elongation and as a docking receptor for adaptor proteins, such as SQSTM1 (also known as p62), that deliver cargo for degradation [21]. Mature autophagosomes, containing organelles and proteins, fuse with lysosomes. The content is degraded and the products are recycled into metabolic and biosynthetic pathways.

In cancer, autophagy can be neutral, tumor-suppressive, or tumor-promoting in different contexts [11]. For example, autophagy can act as a tumor-suppressive mechanism during the early stages of tumorigenesis by suppressing reactive oxygen species (ROS), DNA damage, tissue damage, inflammation, and genome instability, which are known inducers of tumor initiation [22]. However, complete autophagy deficiency in mouse models, even though increasing tumor initiation, is incompatible with tumor progression to a malignant stage, indicating that many cancers benefit from and require functional autophagy for their growth and progression [22]. Moreover, some of the same autophagy functions that initially suppress tumor development render late-stage tumor cells more stress-tolerant and confer resistance to cancer drugs [23]. The mechanism by which autophagy allows resistance to therapy still remains unknown and is likely to be multifaceted, probably involving its central role in metabolic homeostasis [9, 23].

The metabolism-protective role of autophagy implies that autophagy defects render tumor cells hypersensitive to metabolic perturbation and open a therapeutic window for metabolic drug treatment. In support of this concept, mTOR upregulation, which is commonly observed as a drug resistance mechanism in a broad range of tumor types [24, 25], blocks the autophagy pathway at various steps [13] and renders tumors cells hypersensitive to well-tolerated doses of metabolic inhibitors that have little to no effect on normal autophagy-competent cells [15]. In principle, autophagy can also be inactivated genetically. While single-nucleotide variants or short insertion– deletion mutations (commonly referred to as ‘mutations’) in ATGs are remarkably rare in human cancer [26], haploinsufficiency network-based analysis has identified autophagy as a pathway that is frequently hit by somatic copy-number alterations (SCNAs), leading more often to losses than gains [27]. SCNAs in ATGs are usually not homozygous losses, but rather monoallelic deletions affecting multiple ATGs in parallel [27]. As single-allele losses can reduce messenger RNA (mRNA) levels [28], monoallelic deletions hitting multiple ATGs in parallel may limit autophagic flux under metabolic stress and cause a therapeutically relevant vulnerability to metabolic cancer drugs. In support of this hypothesis, RNAi knock-down of BECN1 or LC3 reduces autophagy flux and sensitizes tumors to autophagy inhibitors such as chloroquine [27]. However, formal proof that the accumulation of monoallelic deletions in the autophagy pathway sensitizes to metabolic cancer drugs has been missing.

Here, we have used CRISPR/Cas9 technology to engineer tumor cells with various deletions of the ATGs which are most frequently hit by SCNAs: BECN1, MAP1LC3B/LC3 and ATG10 [27]. While homozygous deletions of every single ATG blocked autophagy and strongly sensitized to inhibitors of aerobic glycolysis, non-homozygous deletions — even when hitting all three ATGs in parallel — did not create a druggable metabolic vulnerability. Together this illustrates that the autophagy pathway of tumor cells is much more robust and resistant to somatic copy-number losses than previously anticipated.

## 2. Materials and Methods

### 2.1. Cell Culture

The NCI-H460 cell line was obtained from the American Tissue Collection Center (ATCC) and cultivated in a humidified atmosphere at 37°C and 5% CO2 using RPMI medium (Gibco) supplemented with 10% fetal bovine serum (FBS, Sigma-Aldrich), 100 U ml^−1^ penicillin, and 100 μg ml^−1^ streptomycin (Life Technologies). Metabolic drugs were obtained from Sigma-Aldrich and used at the following concentrations: dichloroacetate (DCA) 20–60 mM, 2-deoxy-D-glucose (2DG) 2.5–40 mM. Standard concentrations for clonogenic growth assays were 40 mM DCA and 10 mM 2DG. H460 cells with hetero- or homozygous *ATG7* inactivation by CRISPR-mediated insertion or frameshift mutations were described previously [15].

### 2.2. CRISPR/Cas9 Gene Editing

Single-guide RNAs (sgRNAs) targeting genes of interest were designed and cloned into the pSpCas9(BB)-2A-Puro vector (pX459) V2.0 (Addgene #62988) using Golden Gate Cloning as described [29]. The following sgRNAs were used: ATG7, AGA AAT AAT GGC GGC AGC TA; LC3sg1, GGG AAG CAC CGT GTT CAT CG; LC3sg5, AGT TGT GAC CTG CTA CAC AT; BECN1sg1, TCC CTG TAA CAA CCC GTA CG; BECN1sg5, GAT CAC ATC ACA TGG TGA CC; BECN1sgExon4A, ATT GAA ACT CCT CGC CAG GA; ATG10sg3, TCC ATC CGT AAG TTT TCA AG; ATG10sg4, CAA GGA GCT CCT GTA GAC TG. Cells were transfected with a single or a pair of pX459 plasmids using Lipofectamine 2000 (Thermo Fisher Scientific) following the manufacturer’s protocol and selected with puromycin (1 μg ml^−1^). Cells were single-cell cloned and analyzed for the presence of gene deletions and mutations by PCR and sequencing.

### 2.3. PCR and Sequencing

Genomic DNA from cell clones was analyzed by PCR using the following primers: LC3_del_FW (1), CAG ACC TCA GTG CCT CGG TCG A; LC3_del_RV (2), AGA GAA CGC CGC AGA TCC AGG T; LC3_int_FW (3), TGG AGG AAG GAC TGG GTC TC; LC3_int_RV (4), GGC TGT CTG GTG ATT CCT GTA A; BECN1_del_FW (1), TCC ACG GCC TCA GGG ATG GAA G; BECN1_del_RV (2), GTG CAC CCC TGG GCA GTT TTC A; BECN1_int_FW (3), TTC ACC ACA TTG GCC GGA CTG C; BECN1_int_RV (4), AGG GCG GAT GTC ACC AAG CTC T; BECN1 ex4 FW (5), AGG GCA TTC TGT CCT CTG CCC C; BECN1 ex4 RV (6), GCC ATG CTG GTC TTC CAC AGG G; ATG10_del_FW (1), TAC AGC ACC TAG AGT CCC GT; ATG10_del_RV (2), GGA AGC AGG GAG AAA AAA TCC TC; ATG10_int_FW (3), ACT CCC TTT TCC TTG CCT CAT AG; ATG10_int_RV (4), CAC GCT GAA GTC TTG ATA CCC T; ATG7ex2_FW (3), AGT TGT GTT TCA AGG TAG CCT G; ATG7ex2_RV (4), CCG TGA GGA TAA CAG AAG ATG ATG. For each gene, one primer pair 1-2 was designed to amplify a fragment of more than 10 kbp in size spanning most of the gene (Figure 1a). Under standard PCR conditions these primers could only amplify alleles containing a large deletion of the intervening sequence. A PCR product with these primers identified clones with at least one deleted allele (Figure 1b, d). Another primer pair 3-4 was designed to amplify a small internal fragment that indicated the presence of a non-deleted allele (Figure 1b, d). In the case of *BECN1* and *ATG10*, which are present in three copies in H460 cells, clones with monoallelic (+/+/–) and biallelic (+/–/–) deletion were distinguished by sequencing the PCR product of primer pair 1-2, which spans the fusion point. The presence of two different fusion sequences identified clones with two deleted alleles. The *BECN1*^+/–/−^ clone was re-transfected with a nuclease targeting exon 4 and re-cloned. PCR amplification of exon 4 (primers 5-6) and sequencing identified two *BECN1*^fs/–/−^ clones with frameshift-inducing base pair insertions, respectively (Figure 1c). Sequencing of PCR amplicons was performed by LGC Genomics GmbH (Berlin, Germany).

**Figure 1.**
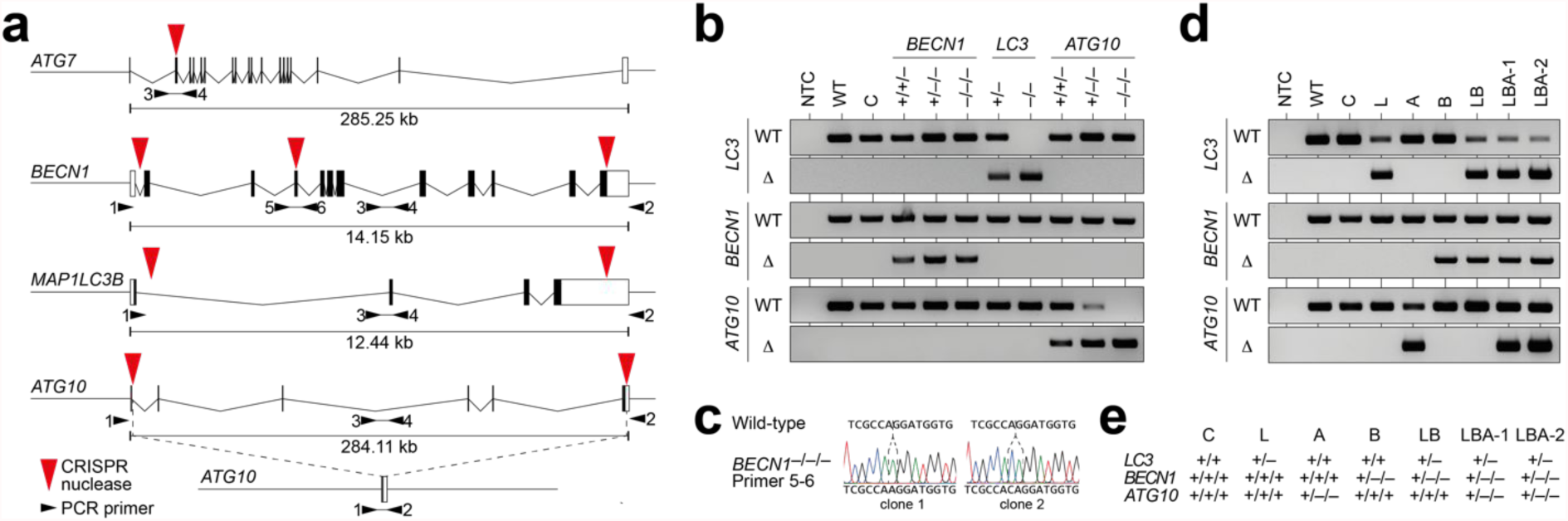
Generation of NSCLC (H460) cells with CRISPR-engineered autophagy gene deletions. (**a**) Genomic structure of the targeted ATGs. Red triangles mark sgRNA target sites, black arrows denote primers used for PCR validation of deletions. Primer pair 1-2 was designed to yield a product only if the intervening sequence between two sgRNA target sites has been deleted (as illustrated for *ATG10* as an example); whereas primer pair 3-4 amplifies the non-deleted alleles. (**b**) PCR validation of clones with deletions of single ATGs. Δ, deleted allele (primer pair 1-2); WT, non-deleted allele (primer pair 3-4). (**c**) Sequencing of *BECN1*^fs/–/−^ reveals frameshift mutations in the non-deleted *BECN1* allele. (**d**) PCR validation of clones with deletions in multiple ATGs. (**e**) Simplified nomenclature of clones with deletions in multiple ATGs. NTC, no template control. C, control clone transfected with non-targeting nuclease; WT, wild-type H460.

### 2.4. Western Blotting

Cells were harvested and lysed with NP-40 Lysis Buffer (50 mM Tris-HCl, 150 mM NaCl, 5 mM ethylenediaminetetraacetic acid (EDTA), 2% NP-40, pH 8.0) supplemented with protease inhibitor (complete ULTRA tablets EASYpack, Roche). Protein yield was quantified via Bradford assay (Bio-Rad). A total of 35-45 μg of proteins were loaded on NuPAGE SDS Gels (Life Technologies) and wet-transferred to PVDF membranes. Membranes were blocked in tris-buffered saline with polysorbate 20 (TBST; 5 mM Tris, 15 mM NaCl, 0.1% Tween 20, pH 7.5) with 5% non-fat dry milk. The primary antibodies used were: LC3B (LC3-I/II) (1:1000, ab48394, Abcam), p62/SQSTM1 (1:1000, P0067, Sigma), ATG7 (D12B11) (1:1000, #8558, Cell Signaling), BECN1 (E8) (1:500, sc-48341, Santa Cruz Biotechnology), ATG10 (EPR4804) (1:1000, ab124711, Abcam), β-Actin (AC-15) (1:10000, ab6276, Abcam). Secondary antibody: goat anti-mouse IgG-HRP (1;5000, #A16084, Thermo Fisher Scientific), donkey anti-rabbit IgG-HRP (1:5000, #NA9340, GE Healthcare). Detection: WesternBright ECL Substrate Sirius kit (Biozym).

### 2.5. Clonogenic Growth Assays

Cells were seeded on cell culture plates, treated after 24h and cultivated for up to 10 days. Cells were fixed in 70% ethanol overnight and stained with Crystal Violet (Sigma-Aldrich) solution diluted 1:20 in 20% ethanol. Colony counting was done manually, taking into consideration only colonies with a cell number ≥ 50.

### 2.6. Real-Time Live Cell Imaging

Live cell imaging was performed by seeding cells in 96-well plates in triplicate for each condition. Drug treatment was started the next day. Cell proliferation was recorded for up to 7 days every 4 hours by the IncuCyte S3 Live-Cell Analysis System (Sartorius) with a phase-contrast microscope at 10X magnification. ImageJ software was used to crop pictures and add scale bars. Confluence analysis was performed with IncuCyte S3 2018A software. Area under the curve (AUC) was calculated with GraphPad Prism 8 using the mean of the first two time points as baseline. AUC values were normalized to the untreated control.

### 2.7. TCGA Pan Cancer Atlas Analysis

TCGA Pan Cancer Atlas data were downloaded from the cBioPortal platform [30]. In our study we included the 9,889 patient samples for which gene copy number and mRNA expression data for *BECN1, ATG7* and *SQSTM1* were available. Log2-transformed copy number data were from the Affymetrix SNP6 platform, mRNA expression data were used as mRNA Z-scores relative to diploid samples (RNA Seq V2 RSEM). The correlation analysis between copy number and protein levels was restricted to 308 samples for which additional protein expression data measured by mass spectrometry were available from the Clinical Proteomic Tumor Analysis Consortium (CPTAC) [31]. All analyses and statistics were calculated with GraphPad Prism 9 Software.

### 2.8. Statistical Analysis

GraphPad Prism 9 Software was used to generate all plots and perform statistical analysis. All data are shown as mean±SD with sample numbers indicated in figure legends. Individual data points are shown as dots. Experiments comparing multiple groups were analyzed by one-way analysis of variance (1way ANOVA) followed by multiple comparison testing according to Dunnett. Experiments investigating the interaction of two variables (e.g. cell type and treatment) were analysed using two-way analysis of variance (2way ANOVA) followed by multiple comparison testing according to Dunnett. P values are reported as follows: ***, P<0.0001; **, P<0.001; *, P<0.01.

## 3. Results

In light of reports that cancer cells frequently harbor several non-homozygous deletions affecting core autophagy genes, we investigated whether somatic copy number losses compromise cancer cell proliferation under metabolic stress induced by experimental compounds that target the aerobic glycolysis (Warburg effect).

### 3.1. Generation of cancer cells with CRISPR-engineered copy number losses of autophagy genes

To investigate the impact of copy number losses in ATGs, we chose the NSCLC cell line NCI-H460 (short H460) as a cellular model. H460 cells are autophagy-competent and respond to inhibitors of aerobic glycolysis with increased autophagic flux [15]. Knock-out of *ATG7* blocks the autophagy response and triggers energy stress and apoptosis, indicating that autophagy induction is essential for H460 cells to maintain metabolic homeostasis under therapeutic perturbation of glycolysis [15]. H460 are hypotriploid and contain 3 copies of *BECN1* and *ATG10* and 2 copies of *MAP1LC3B/LC3* and *ATG7*. All these characteristics together make H460 a valuable model to study how multiple copy number losses of ATGs impact on tumor cell proliferation under metabolic stress.

To delete one or more copies of single ATGs with CRISPR/Cas9 technology, we have generated nuclease pairs which delete the entire coding region of these genes (Figure 1a). Following transfection of Cas9-sgRNA expression plasmids, single-cell clones were isolated and tested by PCR and Sanger sequencing for the presence of deleted and residual wild-type alleles (Figure 1b-e). We obtained *BECN1*^+/+/−^, *BECN1*^+/–/−^ (B), *ATG10*^+/+/−^, *ATG10*^+/–/−^ (A), *ATG10*^−/–/−^ (*ATG*^KO^), *LC3*^+/−^ (L), and *LC3*^−/−^ (*LC3*^KO^) cells (Figure 1b). As this approach failed to generate cells with a deletion of all *BECN1* alleles, we re-transfected *BECN1*^+/–/−^ cells with a single nuclease to introduce frameshift (fs) mutations into exon 4 of the remaining non-deleted allele, yielding two genetically distinct *BECN1*^fs/–/−^ (*BECN1*^KO^-1/2) cell clones (Figure 1c). *ATG7*^+/−^ and *ATG7*^−/−^ (*ATG7*^KO^) cells have been described previously and were used as a reference [15]. Cells transfected with a non-targeting nuclease were used as control (C).

To engineer compound deletions in multiple ATGs, the *LC3*^+/−^ cells were re-targeted sequentially with sgRNA-pairs for *BECN1* and *ATG10* yielding *LC3*^+/−^; *BECN1*^+/–/−^ (short LB) and *LC3*^+/−^; *BECN1*^+/–/−^*;ATG10*^+/–/−^ (short LBA) cells with single remaining copies of the two or three genes, respectively (Figure 1d-e).

### 3.2. BECN1, LC3 and ATG10 knock-out induce hypersensitivity to 2DG and DCA

We first investigated whether *BECN1, LC3* and *ATG10* are essential for maintaining tumor cell proliferation under metabolic stress, similar as has been reported for *ATG7* [15]. Western blots of all knock-out clones confirmed the lack of the targeted proteins (Figure 2a). Consistent with the critical enzymatic role of ATG7 and ATG10 in LC3 lipidation, *AT7*^KO^ and *ATG10*^KO^ cells showed an LC3-I but no (lipidated) LC3-II band, indicative of impaired LC3-I to LC3-II conversion. In *BECN1*^KO^ cells, LC3 conversion was still possible, but resulted in an accumulation of LC3-II. All knock-out clones showed a more or less pronounced accumulation of SQSTM1/p62 (short p62). LC3-II and p62 are both degraded by autophagolysosomes and therefore serve as common markers to estimate autophagic flux [32]. Accumulation of p62 (and LC3-II in *BECN1*^KO^ cells) is therefore consistent with impaired p62/LC3-II degradation due to an autophagy defect.

**Figure 2.**
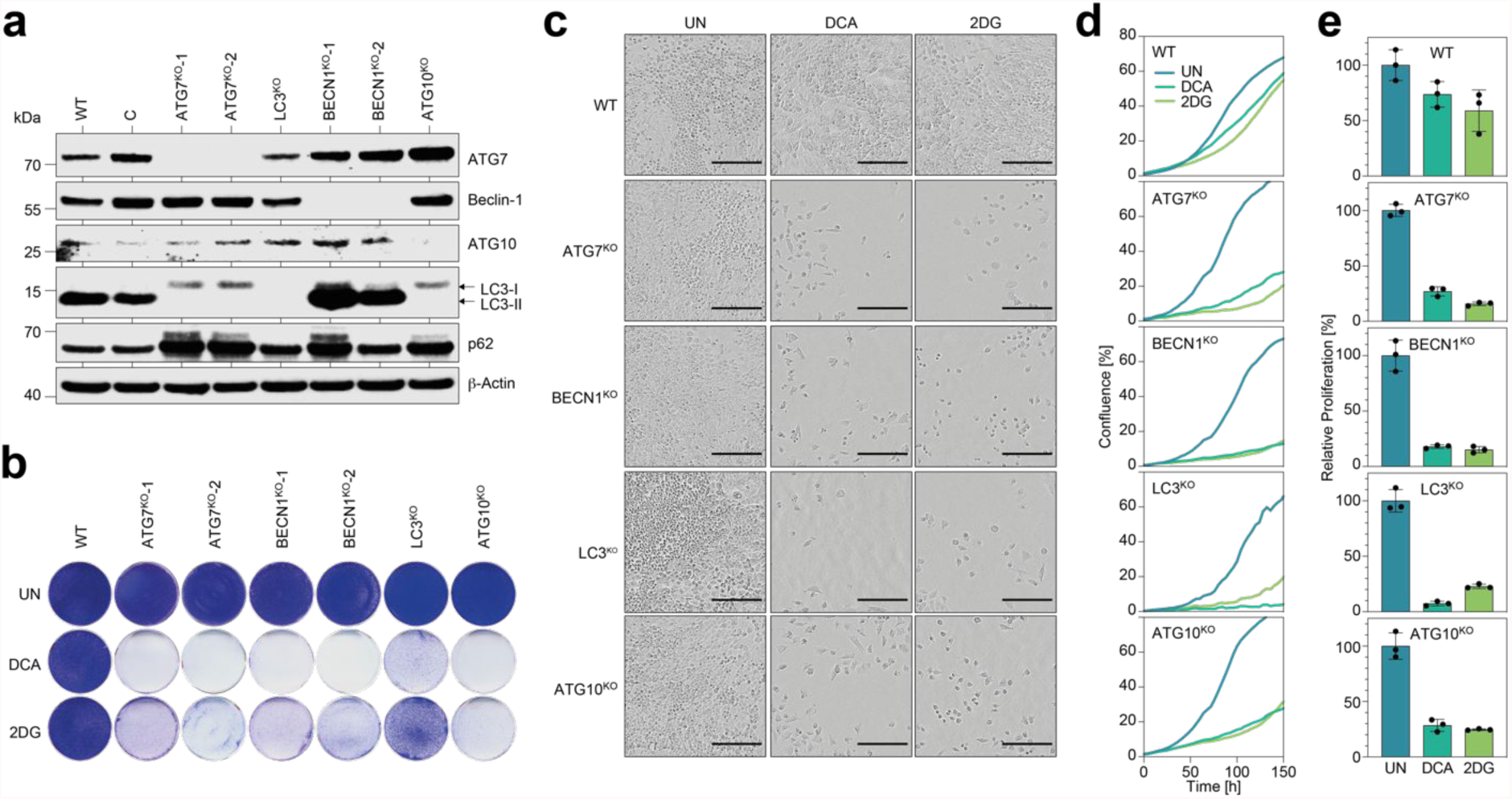
Knock-out of *BECN1, LC3, ATG10* or *ATG7* induces hypersensitivity to 2DG and DCA. (**a**) Western Blot of indicated cells for autophagy genes. β-Actin is shown as loading control; C, control clone transfected with non-targeting nuclease; WT, wild-type H460. (**b**) Clonogenic growth assay. (**c-e**) Real-time live cell imaging of cells under treatment. (**c**) Morphology. Scale bar = 200 μm. (**d**) Proliferation kinetics. Shown is the mean confluence in % (n=3). (**e**) Proliferation. Shown is the area under the curve (AUC) from the curves in (**d**) relative to untreated; mean±SD (n=3). UN, untreated.

We next investigated whether the complete knock-out of any one of the ATGs compromises clonogenic growth under metabolic drug treatment. As metabolic drugs, we used 2DG and DCA which inhibit aerobic glycolysis in a mechanistically distinct manner. Whereas 2DG blocks influx of glucose into glycolysis at the initial step catalyzed by hexokinase, DCA inhibits pyruvate dehydrogenase kinase (PDK), which stimulates efflux from glycolysis into the TCA cycle and thereby decreases flux of glycolytic intermediates into biosynthetic pathways. Clonogenic growth under 2DG or DCA treatment was strongly reduced by knock-out of either ATG (Figure 2b). Time course analysis of cell proliferation by real-time live cell imaging demonstrated that none of the autophagy gene knock-outs impacted proliferation under standard cell culture conditions. In parental cells, proliferation, measured as the area under the confluence curve, decreased under 2DG and DCA treatment by an average of 1.5-fold. In contrast, all knock-out cells showed a severe (3.8-6.0-fold) decrease in proliferation under 2DG and DCA treatment (Figure 2d-f).

We conclude, that tumor cell proliferation under metabolic drug treatment is not only dependent on *ATG7*, but to an equal extent on *BECN1, LC3* and *ATG10*.

### 3.3. Tumor cells with copy number losses in single autophagy genes maintain proliferation under metabolic stress

As autophagy genes are not frequently affected by homozygous ATG deletions, we next explored the functional consequences of non-homozygous deletions (copy number losses) of individual autophagy genes using the CRISPR-engineered cell clones depicted in Figure 1. We first analyzed the impact of copy number losses on protein expression (Figure 3a).

**Figure 3.**
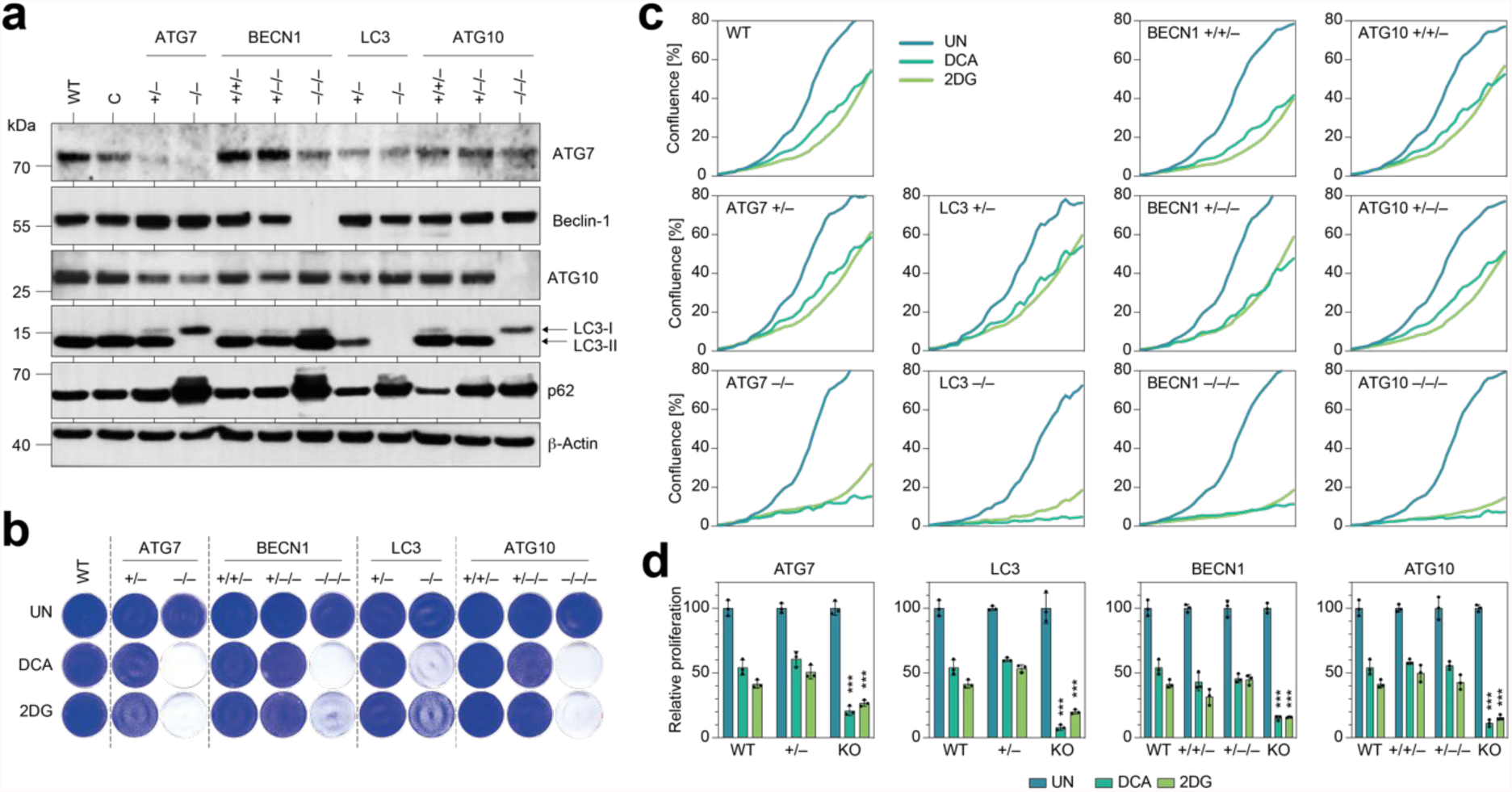
Tumor cells with non-homozygous copy number losses in single ATGs maintain proliferation under metabolic stress. (**a**) Western Blot for ATGs. β-Actin is shown as loading control; C, control clone transfected with non-targeting nuclease; WT, wild-type H460. (**b**) Clonogenic growth assay. (**c-d**) Real-time live cell imaging of cells under treatment. (**c**) Proliferation kinetics. Shown is the mean confluence in % (n=3). (**d**) AUC of proliferation curves shown in (**c**) relative to untreated; mean±SD (n=3). Differential treatment responses were tested for statistical significance by 2way ANOVA and Dunnett’s multiple comparisons test. P-values denote pair-wise comparisons of mutant with parental wild-type cells. UN, untreated.

While the complete knock-out resulted in loss of protein expression, as already seen in Figure 2, non-homozygous copy number losses reduced protein levels of ATG7, Beclin-1, ATG10 and LC3 only weakly (Figure 3a), indicating that even a single remaining allele is sufficient to maintain close to wild-type levels of protein expression. Consistent with the modest effect on protein levels, we only observed mild or no LC3 processing defects and little accumulation of p62 in cells with a single remaining copy of *ATG7, BECN1, LC3* and *ATG10*. In assays for clonogenic growth, only knock-out cells – but none of the cells with non-homozygous copy number losses – showed impaired proliferation under 2DG/DCA (Figure 3b). Consistently, cells with a single remaining copy of either *ATG7, BECN1, LC3* or *ATG10* proliferated under 2DG/DCA treatment with no significant differences to wild-type cells and failed to show any signs of significant hypersensitivity (Figure 3c-d).

We conclude, that single copies of the *ATG7, BECN1, LC3* and *ATG10* gene are largely sufficient to sustain autophagy and maintain tumor cell proliferation under treatment with 2DG and DCA.

### 3.4. Tumor cells with copy number losses in multiple autophagy genes maintain proliferation under metabolic stress

We next explored the consequences of simultaneous non-homozygous copy number losses in multiple ATGs using the engineered cell clones LB and LBA-1/2 (Figure 1 and 4).

**Figure 4.**
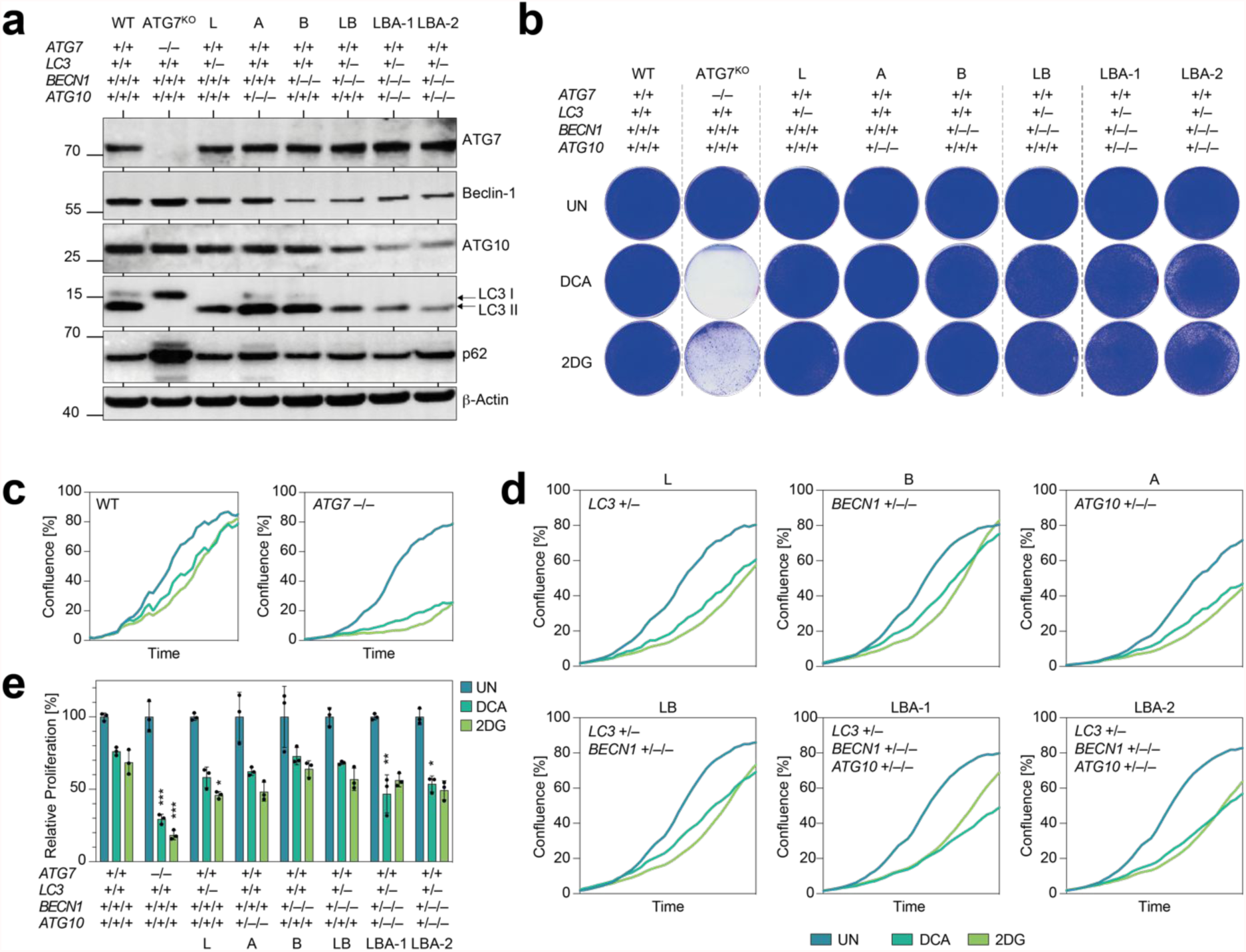
Tumor cells with copy number losses in multiple autophagy genes maintain proliferation under metabolic stress. (**a**) Western Blot of indicated cells for autophagy genes. β-Actin is shown as loading control. (**b**) Clonogenic growth assay. (**c-e**) Real-time live cell imaging of cells under treatment. (**c**,**d**) Proliferation curves. Shown is the mean confluence in % over time (n=3). (**e**) AUC of proliferation curves in (**c**,**d**) relative to untreated; mean±SD (n=3). Differential treatment responses were tested for statistical significance by 2way ANOVA and Dunnett’s multiple comparisons test. P-values denote pair-wise comparisons of mutant with parental wild-type cells. UN, untreated.

Again, we first analyzed the impact of copy number losses on protein expression (Figure 4a). Single-copy expression of *LC3* and *BECN1* (*LC3*^+/−^ *BECN1*^+/–/−^; LB clone) caused modest reduction in Beclin-1 and LC3 protein levels. Similarly, single-copy expression of *LC3, BECN1* and *ATG10* (*LC3*^+/−^ *BECN1*^+/–/−^ *ATG10*^+/–/−^; LBA clones) reduced protein expression of all three proteins. Nevertheless, LC3 conversion defects resulting in LC3-I accumulation were not detected. Moreover, neither LB nor LBA clones consistently showed accumulation of p62 to levels approaching those in single-gene knock-outs (Figures 2a, 3a, 4a). These results suggest that the observed reduction in autophagy protein expression did not reduce autophagic flux under basal growth conditions. In assays for clonogenic growth, only the two LBA clones showed a faint decrease in staining, which by far did not come close to the decrease observed with *ATG7* knock-out cells (Figure 4b). Although statistically significant decreases in proliferation were observed for some of the non-homozygous deletion clones under either DCA or 2DG treatment, proliferation was overall only mildly reduced and did not consistently decrease further with more deletions (Figure 4c-d). Most importantly, none of the non-homozygous deletion clones showed an anti-proliferative DCA/2DG response approaching that of homozygous knock-out clones.

To better identify small anti-proliferative effects of non-homozygous gene deletions, we also conducted the clonogenic growth assay in a quantitative manner. We seeded cells at much lower (clonal) density, treated with 2DG or DCA and counted the emerging colonies manually (Figures 5a-b). Clonogenic growth of B (*BECN1*^+/–/−^), LB (*LC3*^+/−^ *BECN1*^+/–/−^) and LBA-1 *LC3*^+/−^ *BECN1*^+/–/−^ *ATG10*^+/–/−^) cells was significantly decreased by 2DG treatment compared to wild-type cells, but the reduction in colony number was mild (1.8-2.6-fold) compared to 1.3-fold in wild-type cells and 240-fold in homozygous *ATG7* knock-out cells. Notably, the reduction in colony number did not consistently decrease with more deletions. For DCA, the reduction in colony number was not significantly different for the non-homozygous deletion clones compared to wild-type cells. Together these results confirm that single-copy expression of these core autophagy genes is largely sufficient to maintain cell proliferation under metabolic drug treatment (Figure 5b).

**Figure 5.**
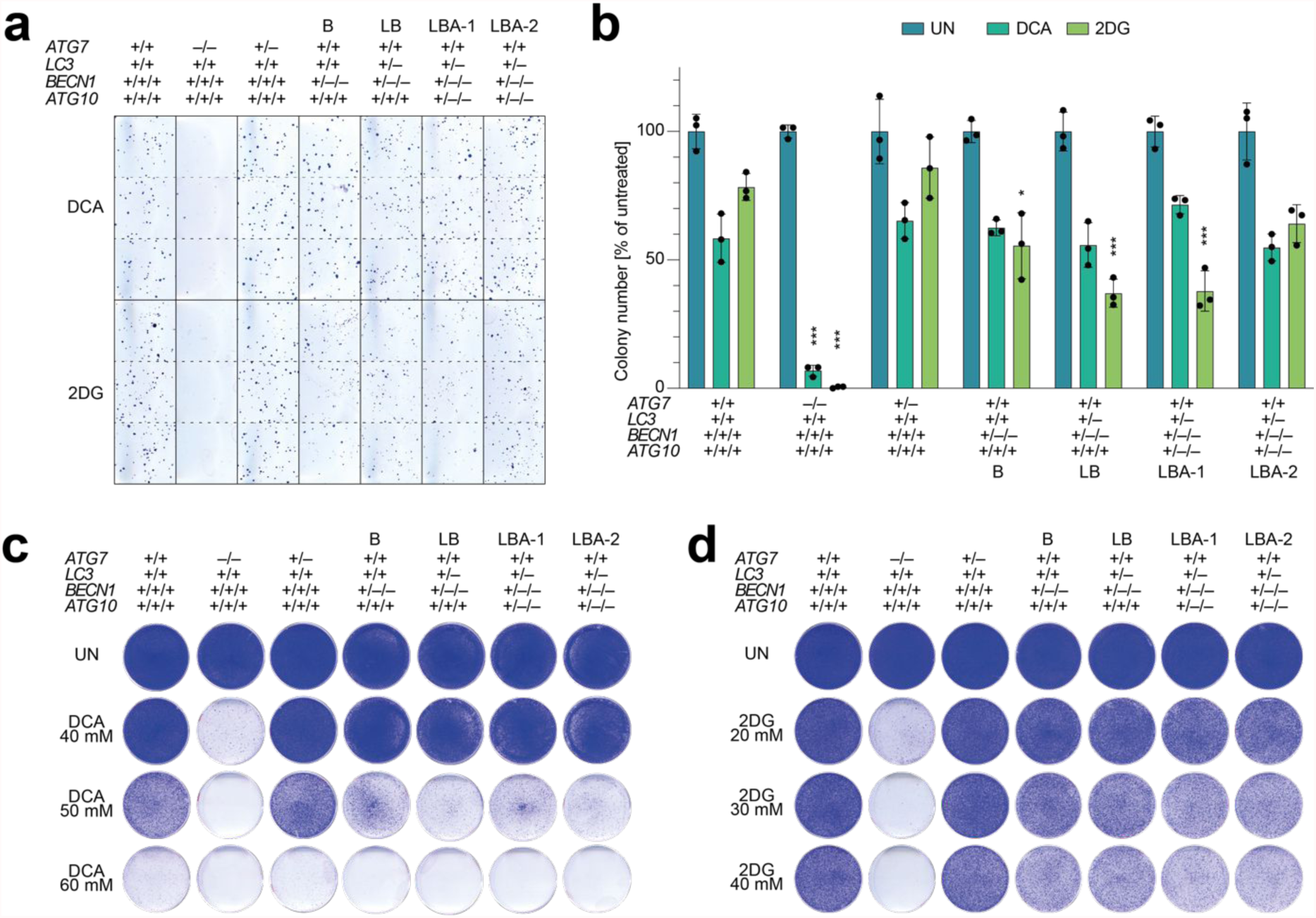
Tumor cells with copy number losses in multiple autophagy genes maintain clonogenic growth under metabolic stress (**a-b**) Quantitative clonogenic growth assay. (**a**) Representative images. (**b**) Colony number in % of untreated cells. Shown is the mean±SD (n=3). Differential treatment responses were tested for statistical significance by 2way ANOVA and Dunnett’s multiple comparisons test. P-values denote pair-wise comparisons of mutant with wild-type cells. (**c-d**) Clonogenic growth under increasing doses of (**c**) DCA and (**d**) 2DG. UN, untreated.

To explore if differential sensitivity becomes apparent at a higher drug concentration, we performed clonogenic growth assays with an extended concentration range (Figures 5c-d). With increasing dose of 2DG and DCA, LB and LBA clones became more sensitive, but this was seen to similar extent also in all other tested cells. Only at an intermediate dose of 50 mM DCA and 2DG doses of 30-40 mM, the B, LB and LBA clones were somewhat more sensitive than wild-type and *ATG7*^+/−^ cells. However, under none of the tested conditions did cells with single-copy expression of *LC3, BECN1* and *ATG10* show a comparable degree of metabolic drug sensitivity as cells with homozygous knock-out of *ATG7*.

### 3.5. Impact of copy number losses in autophagy genes pan-cancer analysis

We finally investigated whether these results can be extrapolated to a broader panel of tumors, represented by the TCGA PanCancer Atlas. We focused our analysis on *BECN1, ATG7* and *SQSTM1/p62* for which data on copy number and both mRNA and protein expression were available. Consistent with an overall close correlation between copy number and mRNA expression [33], copy number of *BECN1* and *ATG7* correlated significantly with mRNA levels across the entire pan-cancer dataset comprising 9,889 patient tumors (Figure 6a and b). In patients, for which mass spectrometry protein data were publicly available from the Clinical Proteomic Tumor Analysis Consortium (CPTAC) [31], *BECN1* and *ATG7* copy number also correlated with the respective protein level (Figure 6c and d). However, correlation at the protein levels was much less obvious than at the mRNA level, indicating that the effect of copy number on protein levels is mitigated by post-transcriptional mechanisms.

**Figure 6.**
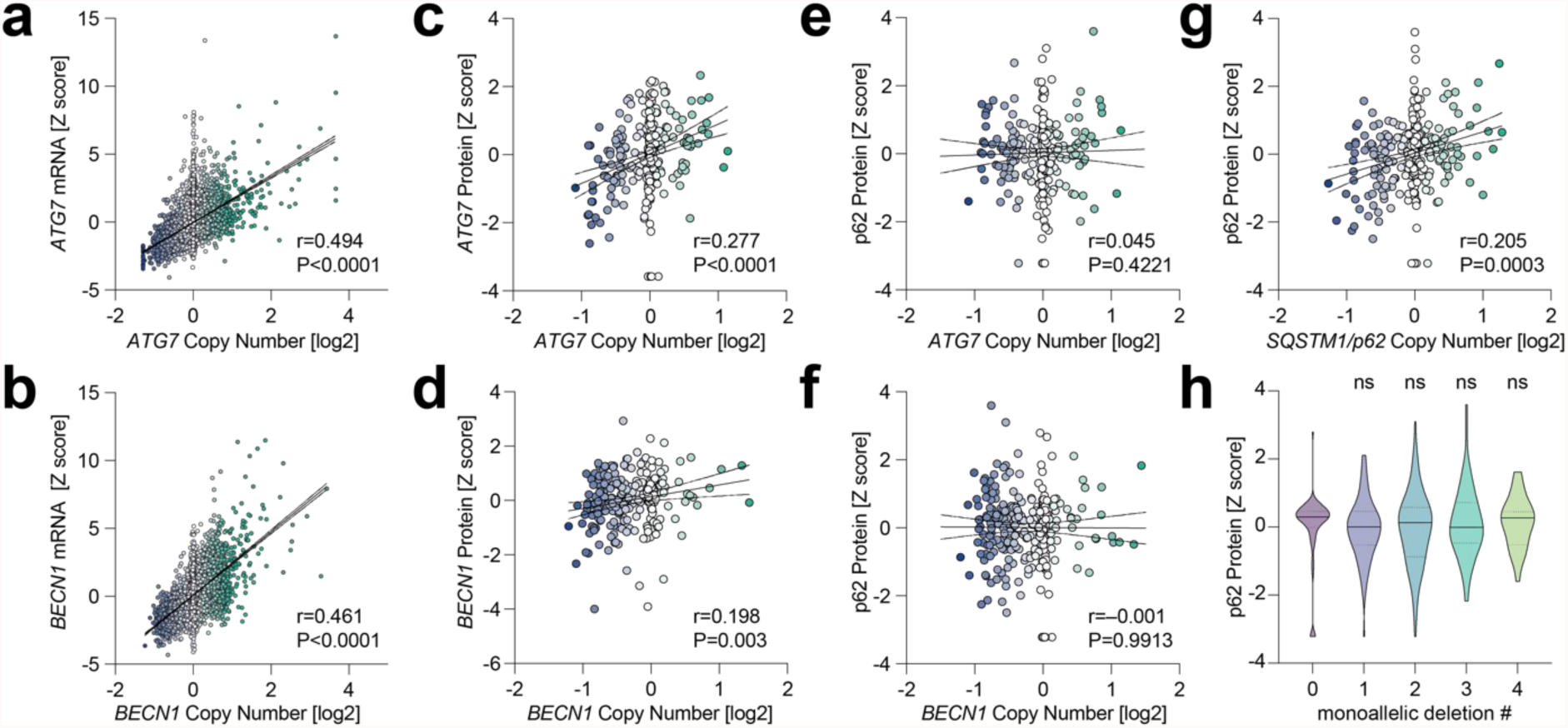
TCGA Pan Cancer Atlas analysis. (**a**,**b**) Correlation analysis between gene copy number (log2-transformed values) and mRNA expression (Z-scores relative to diploid samples). Shown are individual tumors (n=9,889), the linear regression line with 95% confidence interval, the Spearman r correlation coefficient and P value. (**c**,**d**) Correlation analysis between gene copy number (log2-transformed values) and protein expression Z-scores measured with mass spectrometry by the Clinical Proteomic Tumor Analysis Consortium (CPTAC). Shown are individual tumors (n=308), the linear regression line with 95% confidence interval, the Spearman r correlation coefficient and P value. (**e**-**g**) Correlation analysis between p62 protein expression Z-scores (CPTAC) and gene copy number for *ATG7, BECN1* and *SQSTM1/p62*. Shown are individual tumors (n=308), the linear regression line with 95% confidence interval, the Spearman r correlation coefficient and P value. (**h**) p62 protein expression Z-score of patient samples (n=308) stratified according to the number of autophagy genes (*BECN1, LC3, ATG10* and ATG7) harboring shallow (non-homozygous) deletions. Violin plots show the median and interquartile range; 1way ANOVA with Dunnett’s multiple comparisons test. Pairwise comparisons to samples with no deletion in the analyzed autophagy genes were not significant (ns).

We also analyzed whether *BECN1* and *ATG7* copy number impacts on autophagic flux, taking p62 protein accumulation as a surrogate marker for autophagy defects (Figures 6e-h). Notably, p62 protein levels significantly correlated with *SQSTM1/p62* copy number (Figure 6g), validating p62 protein data quality. However, p62 protein did not change in response to varying *BECN1* and *ATG7* copy number (Figures 6e and f). We also assessed whether p62 protein levels might be altered when more ATGs are deleted in parallel, focusing on the genes that we analyzed in our cell line study (*BECN1, ATG7, ATG10, LC3*). Interestingly, p62 protein levels were not correlated with the number of ATGs harboring shallow (non-homozygous) deletions (Figure 6h). p62 levels were not even significantly increased in tumors with shallow deletions in all 4 genes compared to tumors without any deletion. Together this study confirms for patient tumor samples, similar to our observations in the cell line model, that the functional consequences of ATG copy number losses are blunted probably at multiple levels so that autophagic flux is not compromised.

## 4. Discussion

Autophagy plays a key role in maintaining metabolic homeostasis during tumor progression and cancer therapy [11, 12]. This pro-survival activity makes autophagy an attractive target for cancer therapy and various lines of evidence confirm that inhibition of autophagy has the potential to eradicate even advanced tumors. For example, acute autophagy ablation in mice with preexisting NSCLC blocked tumor growth, promoted tumor cell death, and generated more benign disease (oncocytomas) [34]. While indicating that tumors are more dependent on autophagy than normal tissues, these studies also demonstrated that systemic autophagy deficiency in adult mice leads to gradual depletion of dedicated nutrient stores, which is further accelerated by fasting and can result in death from hypoglycemia [34]. To avoid toxicities of systemic autophagy inhibition, an alternative therapy strategy would be to identify tumors with intrinsic autophagy defects that render the metabolism of such tumor cells more susceptible to perturbation and treat those tumors with metabolic inhibitors.

An autophagy defect can be caused by negative regulators of autophagy such as mTOR [13, 35]. mTOR signaling is frequently upregulated in tumors with acquired cancer drug resistance as it, for example, protects from the cytotoxic DNA crosslinks induced by cisplatin by promoting DNA repair and counteracts the cytotoxic activity of PI3K-inhibitors (PI3Ki) by blocking apoptosis [24, 25]. However, as a collateral damage, mTOR blocks autophagy and thereby simultaneously sensitizes such cisplatin- or PI3Ki-resistant tumor cells to drugs that interfere with glycolysis, i.e. the Warburg effect [15]. Importantly, drugs that target glycolysis also affect normal cellular metabolism and are therefore limited more by toxicities than by their ability to kill cancer cells [7]. mTOR-mediated inhibition of autophagy, however, sensitized tumors cells to such an extent that tumors could be eradicated in mice using well-tolerated doses of glycolysis inhibitors, arguing that autophagy defects in tumor cells can open a therapeutic window for metabolic cancer treatments.

In principle, the autophagy pathway can also be compromised by genetic alterations. However, despite clear evidence that autophagy suppresses cancer initiation in mouse models, ATGs are not commonly inactivated by homozygous mutations or gene deletions [26], likely because of their essential role for maintaining metabolic homeostasis during tumor progression and cancer therapy [11, 12]. While homozygous deletions are therefore disfavored during tumor evolution, non-homozygous deletions resulting from somatic copy number losses are conceivable and might, especially when affecting multiple ATGs in parallel, compromise autophagic capacity to an extent that is compatible with normal tumor progression but might create a vulnerability to metabolic perturbation. Indeed, the autophagy pathway is frequently affected by somatic copy number alterations (SCNA), with losses being much more common than gains in a broad range of tumors [27]. These include tumor entities that are particularly rich in SCNAs such as ovarian carcinomas, but also in tumors like non-small cell lung cancer (NSCLC), where SCNAs are less dominant drivers of tumorigenesis than point mutations [27, 36]. In some tumors, non-homozygous SCNAs indeed affect multiple ATGs, suggesting that the combined loss synergistically limits autophagic flux especially under autophagy-stressing conditions [27]. In fact, the ovarian cancer cell line OVCAR-3 with monoallelic losses of *BECN1* and *LC3* is more sensitive to various autophagy-stressing drugs than the SKOV-3 ovarian cancer cell line without these deletions, while the latter is sensitized by RNAi knock-down of either BECN1 or LC3 [27]. Together, these results strongly suggested that monoallelic losses – especially when affecting multiple genes at once – confer a druggable vulnerability. We therefore hypothesized that ATG copy number losses might identify tumors which are also less capable of compensating inhibition of aerobic glycolysis through autophagy.

Although the NSCLC cells in our study were strongly sensitized to 2DG and DCA by knock-out of single ATGs (Figure 2), we did not observe a similar metabolic vulnerability when we experimentally deleted only single copies with CRISPR nucleases (Figure 3). Importantly, we did not even observe a comparable vulnerability when deleting three core autophagy genes down to a single remaining copy of each (Figure 4 and 5). Notably, we observed only a modest reduction of protein levels and weak signs of decreased autophagic flux such as reduced LC3 conversion and p62 accumulation. In general, SCNAs are well-known to alter the expression level of genes located at the affected genomic region and thereby contribute to or even drive tumor development [37-39]. There is an overall close correlation between gene copy number variation and differential gene expression across a broad range of cancer types and the copy number displays a positive linear influence on gene expression for the majority of genes, indicating that genetic variation has a direct effect on gene transcription [33]. Due to transcriptional adaptive mechanisms, however, changes in gene copy number at the genomic level do not always translate proportionally into altered gene expression levels [40-42]. Although most of the autophagy genes do not show significant changes in mRNA levels when autophagy is induced, expression of these genes is nevertheless tightly regulated at the level of transcription and post-transcriptionally, for example, via miRNAs [32, 43, 44]. The mechanisms underlying transcriptional and post-transcriptional adaptation of genes to SCNAs still remain poorly understood and appear to be pathway-dependent as some biological processes show higher degrees of adaptation than others [40, 41, 45]. For the autophagy pathway, wide-spread adaption to SCNAs is supported by our analysis of the TCGA Pan Cancer Atlas cohort (Figure 6). Across all patients, the copy number of key autophagy genes *BECN1* and *ATG7* correlated well with mRNA expression but to a much lesser degree with protein levels (Figures 6a-d). Moreover, although p62 protein correlated with *SQSTM1/p62* copy number, no accumulation of p62 protein was detected in patient samples with shallow deletions, i.e. non-homozygous copy number losses, in single or multiple autophagy genes (Figures 6e-h).

This demonstrates, that the extent of copy number reduction detected at the genome level is strongly decreased at the protein level, likely as a result of both transcriptional and post-transcriptional adaptation, which blunts potential negative effects on autophagic flux that would impede tumor progression or sensitize to cancer therapy.

## 5. Conclusions

Autophagy is essential for metabolic homeostasis during tumor progression and contributes to therapy resistance [11, 12]. Cancer cells, nevertheless, often harbor non-homozygous copy number losses of multiple core autophagy genes [27], suggesting that such tumors are more susceptible to therapy, in particular with compounds that perturb cancer cell metabolism. Using isogenic cell clones with defined CRISPR-engineered autophagy gene deletions, we show that tumor cell proliferation and clonogenic growth can be largely maintained under metabolic perturbation by single-copy expression of core autophagy genes. Only homozygous deletions interfere with tumor cell proliferation under metabolic drug treatment. In patient tumors, non-homozygous autophagy gene deletions result in only little reduction in the respective protein levels and fail to induce p62 accumulation as a sign of reduced autophagic flux. We conclude, that the autophagy pathway is markedly robust to non-homozygous copy number losses – even when multiple genes are affected simultaneously.

## Author Contributions

Conceptualization, T.S. and M.W.; generation and initial characterization of cell clones, P.P.; validation of clone genotype, N.G.; clonogenic growth assays, live cell imaging and Western blots, M.W.; formal analysis, P.P., T.S. and M.W.; resources, T.S.; data curation, P.P., T.S. and M.W.; writing—original draft preparation, P.P. and T.S..; writing—review and editing, all; visualization, all; supervision, M.W..; project administration, M.W..; funding acquisition, M.W.. All authors have read and agreed to the published version of the manuscript.

## Acknowledgments

We acknowledge Aaron Dort and Bjorn Geißert for technical assistance and thank members of the lab for helpful discussion and advice.

